# BCFtools/csq: Haplotype-aware variant consequences

**DOI:** 10.1101/090811

**Authors:** Petr Danecek, Shane A. McCarthy

## Abstract

**Motivation:** Prediction of functional variant consequences is an important part of sequencing pipelines, allowing the categorization and prioritization of genetic variants for follow up analysis. However, current predictors analyze variants as isolated events, which can lead to incorrect predictions when adjacent variants alter the same codon, or when a frame-shifting indel is followed by a frame-restoring indel. Exploiting known haplotype information when making consequence predictions can resolve these issues.

**Results:** BCFtools/csq is a fast program for haplotype-aware consequence calling which can take into account known phase. Consequence predictions are changed for 501 of 5019 compound variants found in the 81.7M variants in the 1000 Genomes Project data, with an average of 139 compound variants per haplotype. Predictions match existing tools when run in localized mode, but the program is an order of magnitude faster and requires an order of magnitude less memory.

**Availability:** The program is freely available for commercial and non-commercial use in the BCFtools package which is available for download from http://samtools.github.io/bcftools

## 1 Introduction

With the rapidly growing number of sequenced exome and whole-genome samples, it is important to be able to quickly sift through the vast amount of data for variants of most interest. A key step in this process is to take sequencing variants and provide functional effect annotations. For clinical, evolutionary and genotype-phenotype studies, accurate prediction of functional consequences can be critical to downstream interpretation. There are several popular existing programs for predicting the effect of variants such as the Ensembl Variant Effect Predictor (VEP) (McLaren *et al.*, 2016), SnpEff (Cingolani *et al.*, 2012) or ANNOVAR (Wang et al., 2010). One significant limitation is that they are single-record based and, as shown in Figure 1, this can lead to incorrect annotation when surrounding in-phase variants are taken into account.

**Fig. 1.**
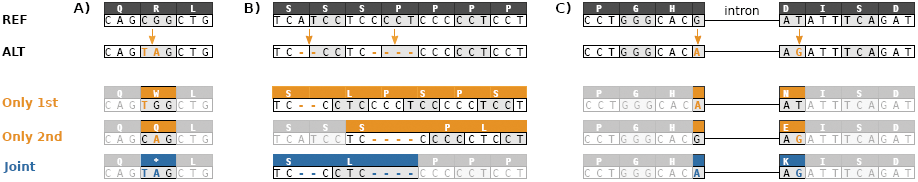
Three types of compound variants that lead to incorrect consequence prediction when handled in a localized manner, i.e. each variant separately rather than jointly. A) Multiple SNVs in the same codon result in a TAG stop codon rather than an amino acid change. B) A deletion locally predicted as frame-shifting is followed by a frame-restoring variant. Two amino acids are deleted and one changed, the functional consequence on protein function is likely much less severe. C) Two SNVs separated by an intron occur within the same codon in the spliced transcript. Unchanged areas are shaded for readability. All three examples were encountered in real data.

With recent experimental and computational advancements, phased haplotypes over tens of kilobases are becoming routinely available through the reduced cost of long-range sequencing technologies (Zheng *et al.*, 2016) and the increased accuracy of statistical phasing algorithms (Sharp *et al.,* 2016; Loh *et al.*, 2016) due to the increased sample cohort sizes (McCarthy *et al.*, 2016). We present a new variant consequence predictor implemented in BCFtools/csq that can exploit this information.

## 2 Methods

For haplotype-aware calling, a phased VCF, a GFF3 file with gene predictions and a reference FASTA file are required. The program begins by parsing gene predictions in the GFF3 file, then streams through the VCF file using a fast region lookup at each site to find overlaps with regions of supported genomic types (exons, CDS, UTRs, or general transcripts). Active transcripts that overlap variants being annotated are maintained on a heap data structure. For each transcript we build a haplotype tree which includes phased genotypes present across all samples. The nodes in this tree correspond to VCF records with as many child nodes as there are alleles. In the worst case scenario of each sample having two unique haplotypes, the number of leaves in the haplotype tree does not grow exponentially but stops at the total number of unique haplotypes present in the samples. Thus each internal node of the tree corresponds to a set of haplotypes with the same prefix and the leaf nodes correspond to a set of haplotypes shared by multiple samples. Once all variants from a transcript are retrieved from the VCF, the consequences are determined on a spliced transcript sequence and reported in the VCF.

Representing the consequences is itself a challenge as there can be many samples in the VCF, each with different haplotypes, thus making the prediction non-local. Moreover, diploid samples have two haplotypes and at each position there can be multiple overlapping transcripts. To represent this rich information and keep the output compact, all unique consequences are recorded in a per-site INFO tag with structure similar to existing annotators. Consequences for each haplotype are recorded in a per-sample FORMAT tag as a bitmask of indexes into the list of consequences recorded in the INFO tag. The bitmask interleaves each haplotype so that when stored in BCF (binary VCF) format, only 8 bits per sample are required for most sites. The bitmask can be translated into a human readable form using the BCFtools/query command. Consequences of compound variants linking multiple sites are reported at one of the sites only with others referencing this record by position.

## 3 Results

### 3.1 Accuracy

Accuracy was tested by running in localized mode and comparing against one of an existing local consequence caller (VEP) using gold-standard segregation-phased NA12878 data (Cleary *et al.*, 2014). With each site treated independently, we expect good agreement with VEP at all sites. Indeed, only 11 out of 1.6M predictions differed within coding regions. See the Supplement (S2) for further details about these differences. Detailed comparison between VEP and other local callers is a topic that has been discussed elsewhere (McCarthy *et al.*, 2014).

### 3.2 Performance

Performance was compared to VEP (McLaren *et al.*, 2016), SnpEff (Cingolani *et al.*, 2012) and ANNOVAR (Wang et al., 2010) running on the same NA12878 data. In localized and haplotype-aware mode, BCFtools/csq was faster by an order of magnitude than the fastest of the programs and required an order of magnitude less memory, see Suppl. Table S1. In contrast to localized calling, scaling of haplotype-aware calling will depend on the number of samples being annotated. In Suppl. Fig. S2-S5, we show that memory and time both scale linearly with number of sites in the transcript buffer and number of samples.

### 3.3 Compound variants in 1000 Genomes

Applied to the 1000 Genomes Phase 3 data, haplotype-aware consequence calling modifies the predictions for 501 of 5019 compound variants, summarised in Table 1 and discussed in the Supplement S3. On average, we observe 139.4 compound variants per haplotype (Suppl. Fig. S1), recover 16.4 variants incorrectly predicted as deleterious, and identify 0.8 newly deleterious compound variants per haplotype.

**Table 1.** Summary of BCFtools/csq consequence type changes from localized (rows) to haplotype-aware (columns) calling in 1000 Genomes data. Blue / orange background indicates a change to a less / more severe prediction in haplotype-aware calling. Only variants with modified predictions are included in the table.

| haplotype aware: |  | synonymous | missense | inframe | stop gained | stop lost |
| --- | --- | --- | --- | --- | --- | --- |
| localized | synonymous | 24 | 5 | 0 | 0 | 1 |
|  | missense | 31 | 4429 | 0 | 66 | 0 |
|  | frameshift | 0 | 0 | 121 | 0 | 0 |
|  | stop gained | 0 | 273 | 1 | 29 | 0 |
|  | stop lost | 0 | 3 | 0 | 0 | 9 |

To highlight an example, a frame-restoring pair of indels in the DNA-binding protein gene SON was found to be monomorphic across all 1000 Genomes samples. There, a 1-bp insertion followed by a 1-bp deletion (G>GA at 21:34948684 and GA>G at 21:34948696) are each predicted as frame-shifting, but in reality the combined effect is a substitution of four amino acids. The functional consequence is therefore likely much less severe, consistent with the SON gene being highly intolerant of loss-of-function mutations, as predicted by ExAC (Lek *et al.*, 2016).

In most studies haplotypes have been determined statistically. Given the typical 1% switch error rate, we estimate the compound error rate from the distribution of heterozygous genotypes in compound variants to be 1.1%, see the Supplement S4 for details.

## 4 Discussion

Correctly classifying the functional consequence of variants in the context of nearby variants in known phase can change the interpretation of their effect. Variants previously flagged as benign or less severe may now be flagged as deleterious and vice versa. In a rare disease sequencing study, for example, this may have a significant impact as these functional annotations may determine which variants to follow up for further study.

Previous work by Wei *et al.* (2015) does not consider indels or introns occurring within the same codon, and requires access to the BAM alignment files to estimate haplotypes. Our approach starts with phased VCF data, leaving haplotype calling as a problem to be solved by other means, for example by statistical phasing. Instead, we focus on providing fast consequence prediction taking into account all variation within a transcript.

The standard programs have rich functionality beyond the reporting of variant consequence, and the aim of BCFtools/csq is not to compete with that. Instead, we propose haplotype-aware calling is included in annotation pipelines for enhanced downstream analysis.

## Acknowledgements

The authors thank Monica Abrudan, Richard Durbin, Daniel Gaffney, Thomas Keane, William McLaren and Kim Wong for helpful discussions and ideas.

### Funding

The work was supported by the Wellcome Trust (WT098051) and a grant co-funded by the Wellcome Trust and Medical Research Council (WT098503).

